# Leftward transfer of a chemosensory polycystin initiates left-dominant calcium signaling for lateralized embryonic development

**DOI:** 10.1101/2023.01.12.523739

**Authors:** Yosuke Tanaka, Ai Morozumi, Nobutaka Hirokawa

**Affiliations:** Department of Cell Biology and Anatomy, Graduate School of Medicine, The University of Tokyo, 7-3-1 Hongo, Bunkyo-ku, Tokyo 113-0033, Japan

## Abstract

Left-dominant [Ca^2+^]_i_ elevation on the margin of ventral node furnishes the initial laterality signaling in mouse embryos. It depends on nodal flow, FGFR signaling, and PKD1L1-containing polycystin channels, of which interrelationship is still missing. Here we show that PKD1L1 protein is predominantly accumulated on the left margin of the nodal pit and serves as a chemosensory channel for Nodal-mediated [Ca^2+^]_i_ elevation. PKD1L1/PKD2 overexpression augmented the Nodal sensitivity of fibroblasts. We detected PKD1L1-containing fragile meshwork of fibrous strands using KikGR-PKD1L1 knockin mice, especially when the extraembryonic membrane was preserved. The portion of meshwork bridging over nodal crown cells significantly lateralized to the left. This bridge was formed by a leftward flow of PKD1L1-containing fibrous strands, which can be suppressed by the FGFR inhibitor SU5402 that antagonized the [Ca^2+^]_i_ elevation as well. These data provide evidence for a leftward transfer of chemosensory PKD1L1 polycystin channel, as a readout mechanism of nodal flow.

## Introduction

The exquisite coordination in body plan formation is promoted by determination of the left-right asymmetry. The mouse left-right organ, ventral node is located at the most posterior part of a concaved ventral midline structure, consisting of columnar epithelium of the notochordal plate being derived from the axial mesoderm (Hamada, 2020; Hirokawa et al., 2006). In the center of this nodal pit, the nodal pit cells (NPCs) mainly generate a leftward fluid flow with their rapidly rotating monocilia (Grimes and Burdine, 2017; Nonaka et al., 1998). As defined previously (Odate et al., 2016), they are surrounded by the nodal crown cells (NCCs) of notochordal plate with less motile cilia, which shape the side wall of the nodal pit. They are further surrounded by the endodermal crown cells (ECCs) that consist of large and flat simple squamous epithelium. The whole structure is enwrapped by an extraembryonic membrane that may preserve the extraembryonic fluids and any fine fragile structures within the nodal pit. We have previously invented a nodal flow hypothesis of lateral symmetry breaking, according to molecular genetics analyses of the cilia-missing *Kif3b*^*-/-*^ mutant mouse embryos with randomized laterality (Nonaka *et al*., 1998). Briefly, posteriorly tilted and clockwise rotating monocilia of NPCs can generate a leftward fluid flow, according to the planar cell polarity (PCP) signaling and a hydrodynamic action from the surface of the NPCs (Hamada and Tam, 2020; Okada et al., 2005). The “two-cilia theory” has been raised by the observation that the cilia on the crown cells are less motile than those on the NPCs, so that the NPC cilia may be mainly involved in flow generation and the crown cell cilia may be mainly involved in a sensory function to evoke the downstream signal transduction on the left side (McGrath and Brueckner, 2003; Tabin and Vogan, 2003).

However, this flow interpretation mechanism on the left side of the ventral node is still controversial. [Ca^2+^]_i_ elevation on the left margin of the nodal pit has been believed to be the closest downstream event of the leftward fluid flow, which may successively evoke the expression of left organizer genes in the left lateral plate mesoderm (McGrath et al., 2003; Mizuno et al., 2020; Tanaka et al., 2005). The Ca channels, PKD1L1/PKD2 (TRPP2) or PKD1L1/PKD2L1 polycystin complexes are thought to be responsible for proper left-right determination via evoking the [Ca^2+^]_i_ elevation, according to genetic evidence in human, mouse, and fish (Field et al., 2011; Grimes et al., 2016; Kamura et al., 2011; Le Fevre et al., 2020; Vogel et al., 2010). They are partly expressed on the cilia in the mouse ventral node or fish Kupffer vesicles (DeCaen et al., 2013; Field *et al*., 2011; Kamura *et al*., 2011; Mick et al., 2015). Cilia are generally involved in chemosensory function in many organisms (Pazour and Witman, 2003; Signor et al., 1999; Singla and Reiter, 2006), and some cilia can also secrete PKD-containing extracellular vesicles (EVs) to facilitate its long-range transfer (Wang et al., 2021; Wood and Rosenbaum, 2015). The idea that these polycystins may serve as a mechanosensor for cilia bending (Djenoune et al., 2023; Katoh et al., 2023; McGrath and Brueckner, 2003; Tabin and Vogan, 2003; Yoshiba et al., 2012) has however been challenged by the experimental evidence for that an artificial fluid flow does not apparently bend the cilia or elevate the intraciliary calcium (Delling et al., 2016; Tajhya and Delling, 2020). In addition, chemosensory properties of the PKD1/2 channels have been identified in other systems (Ha et al., 2020; Horio et al., 2011; Kim et al., 2016). Thus, a similar chemosensory property of PKD channels is also expected in the ventral node, but its ligand has not been determined yet. In addition, the laterality of polycystin protein expression has not been precisely investigated.

In our previous study, left-dominant [Ca^2+^]_i_ elevation in 2–3-somite stage embryos was completely eliminated after treatment by the FGFR inhibitor SU5402 for 1 h, leaving the nodal flow intact. A part of [Ca^2+^]_i_ elevation can be restored by the addition of Shh-N polypeptide (Tanaka *et al*., 2005). The release of huge extracellular materials, which were termed nodal vesicular parcels (NVPs), well correlated with those pharmacological characteristics in [Ca^2+^]_i_ elevation (Tanaka *et al*., 2005). Therefore, it is likely that an FGFR-dependent cellular event must be involved in [Ca^2+^]_i_ elevation on the left side, and that leftward material transfer by the nodal flow is one of its possible mechanisms.

To investigate in vivo dynamics of polycystin, we have knocked in a humanized cDNA for the photoconvertible Kikume Green-Red (KikGR) fluorescent protein tag (Tsutsui et al., 2005) into the N-terminus of mouse *Pkd1l1* gene, by using CRISPR/Cas9 technology. Microscopical observations have revealed a leftward shift of PKD1L1 localization from the midline, which then forms physically robust bridges between NPCs and the left margin of the nodal pit, over the NCCs in an FGFR-dependent manner. PKD1L1 preferentially binds to Nodal on this left margin, which may evoke the left-dominant [Ca^2+^]_i_ elevation as evidenced by in vitro calcium imaging. Those experimental data convey the insight into that the PKD receptor channel itself, rather than its ligands, is an essential effector of the nodal flow that is accumulated on the left side. These findings will unveil one of the most important missing links in the left-right determination pathway.

## Results

### Spatial discrepancy between transcript and protein in *Pkd1l1* gene products

To investigate the spatiotemporal expression pattern of PKD1L1 polycystin, we conducted whole mount in situ hybridization of 7.5 dpc mouse embryos, using a well-characterized *Pkd1l1* riboprobe (Tamplin et al., 2011). The nodal pit of 2–3 somite embryos were predominantly labelled, in addition to a midline structure in more anterior regions (**Fig. 1A and B**).

However, whole mount immunohistochemistry of mouse embryo tail buds after 2 somite stage revealed a left-dominant expression pattern of PKD1L1 protein within and around the nodal pit (**Fig. 1C and D**). Despite the disturbance due to opening the extraembryonic membrane before fixation, left margin of the nodal pit was predominantly labelled by the PKD1L1 antibody. Furthermore, many fluorescently labelled linear arrays appeared to be radiated from there, and observed over the left ECCs. A part of left NCCs were covered by thick bundles of linear arrays like a bridge between the NPCs and the left margin (**Fig. 1C**, between the arrowheads). The existence of those “NCC bridges” were significantly lateralized to the left side (**Fig. 1D**). Because the *Pkd1l1* mRNA expression was limited on the midline (**Fig. 1A**), these lateralized distribution of the PKD1L1 protein must be derived from a posttranslational physical event.

**Figure 1.**
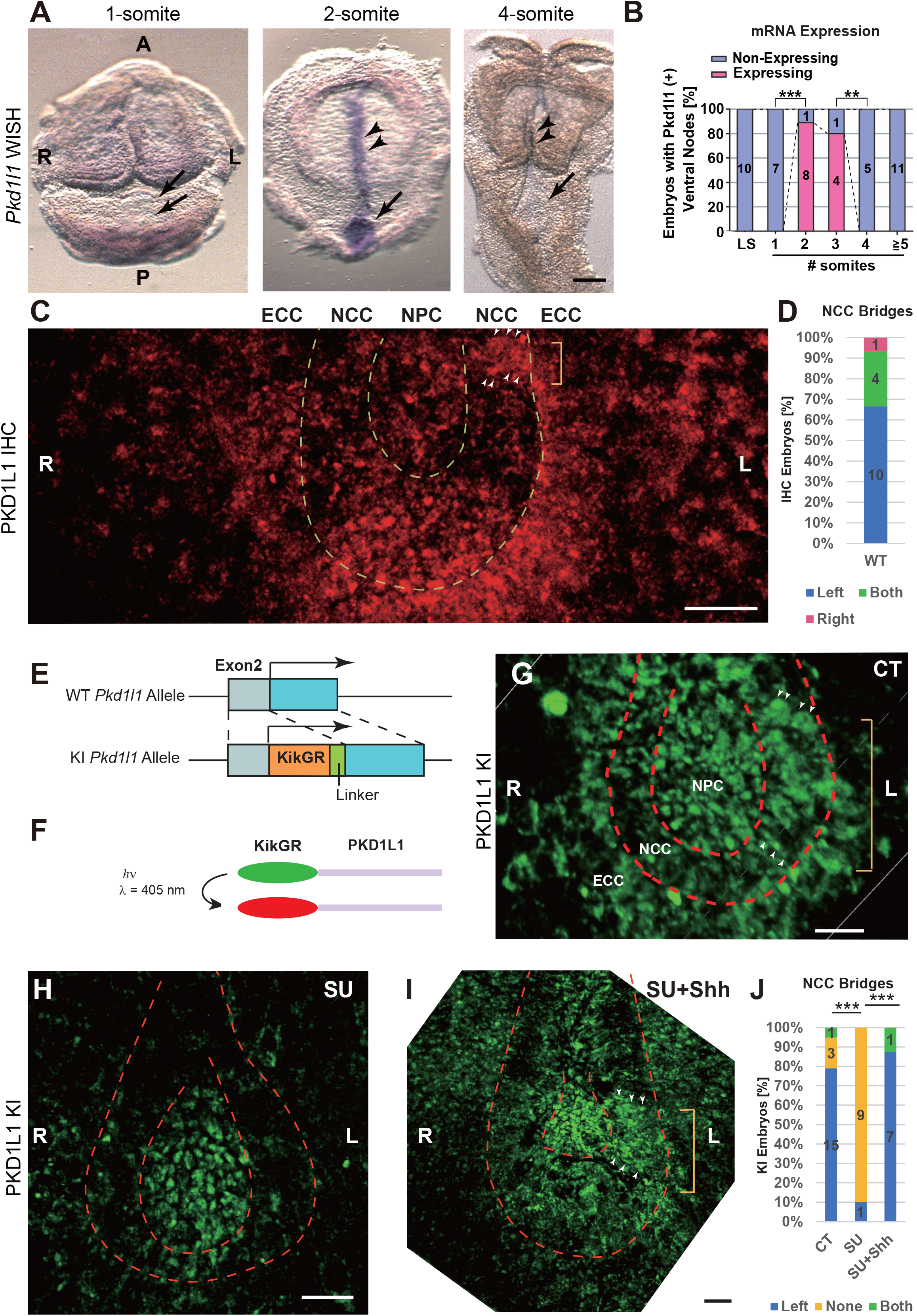
Lateralization of PKD1L1 protein in mouse embryos. (A and B) *Pkd1l1* whole mount in situ hybridization of wild-type mouse embryos at 7.5 dpc (A) and its quantification (B). Scale bar, 100 μm. Arrows, ventral nodes; Arrowheads, notochords; L, left; R, right; A, anterior; P, posterior; throughout the panels. The embryo stages are indicated above the panels. \*\**p* < 0.01; \*\*\**p* < 0.001; One-sided Fisher’s exact test with the sample numbers indicated in the graph. (C and D) *z-*projected PKD1L1 whole mount immunocytochemistry of a 4-somite stage wild type mouse embryo (C), accompanied by statistics of the fraction of embryos of 2– 4-somite stages positive for NCC bridges on the indicated sides (D). ECC, endodermal crown cell; NCC, nodal crown cell; NPC, nodal pit cell; throughout the panels. Dotted lines, margins of the NPC and NCC areas, respectively. The structure between the arrowheads, indicated by the yellow bracket, represents the left NCC bridge. Scale bar, 20 μm. (E and F) Schematic representation of the *KikGR* cDNA knockin strategy into the mouse *Pkd1l1* locus (KI; E), and that of KikGR-PKD1L1 protein photoconversion in living mouse embryos (F). Light blue, 5’-noncoding region. Blue, coding region. Corresponding to **Supplementary Figs. S1 and S2 and Tables S1–S3**. (G–J) *z*-projections of fluorescent ventral views around the ventral nodes of *Pkd1l1*^*KikGR/KikGR*^ embryos, which were preincubated for 1 h in DR75 medium without (CT; G) or with 20 μM SU5402 (SU; H) and/or 2 μg/ml Shh-N as indicated (SU + Shh; I), accompanied by statistics of the NCC bridge laterality (J). Arrowheads in (G) and (I), the margins of left NCC bridges that are indicated by yellow brackets. Scale bars, 20 μm. \*\*\**p* < 0.001, Fisher’s exact test between the left and others. Dotted lines, margins of the NPC and NCC areas, respectively. Corresponding to **Supplementary Fig. S3 and Movie S1**.

### Knockin embryos reproduced PKD1L1-containing NCC bridges on the left

Using the CRISPR/Cas9 technology, we knocked in a *KikGR* cDNA encoding the Kikume Green-Red photoconvertible fluorescent protein (Tsutsui *et al*., 2005), before the start codon of the *Pkd1l1* gene of C57BL/6 mice (**Fig. 1E**; **Supplementary Figs. S1 and S2; Supplementary Tables S1–S3)**. Because KikGR protein can be photoconverted from green to red by the 405-nm light (**Fig. 1F**), it was expected to serve for simultaneously chasing the expression and dynamics of the PKD1L1 protein.

The homozygous mice were healthy and viable without any apparent developmental abnormalities including heterotaxia, suggesting that this N-terminal tagging did not largely affect the protein function. Then, we observed the ventral nodes of the *Pkd1l1*^*KikGR/KikGR*^ mouse embryos by inverted confocal microscopy, after culturing them for 1 h. The green fluorescence on the surface of nodal pit was well colocalized to that of PKD1L1 immunofluorescence in semi-superresolution (**Supplementary Fig. S3A–F**). The green signal intensities in *Pkd1l1*^*KikGR/KikGR*^ mouse embryos were stronger than those in wild type (WT; **Supplementary Fig. S3G and H**). Thus, those signals may largely represent the intrinsic PKD1L1 protein localization. We detected many linear arrays of signal over the left-half of the nodal pit, in embryos later than 2-somite stage (**Fig. 1G** and **Supplementary Fig. S3I**). Intriguingly, the NCC bridge (**Fig. 1G**, the signal between the arrowheads) was again labelled, which was significantly lateralized to the left side (15/19; **Fig. 1J**; **Supplementary Movie S1**).

We further investigated the FGFR/Shh-dependency of this NCC bridge formation (**Fig. 1H–J**). Application of the FGFR inhibitor SU5402 at 20 μM for 1 h resulted in a significant decrease in NCC bridge formation on either side (9/10; **Fig. 1H and J**). Because the green signal in the NPC region was largely preserved after SU5402 treatment, physical properties of PKD1L1-carrying structure may be altered by the FGFR inhibition. Application of SU5402 plus recombinant mouse Shh-N polypeptide significantly reversed this elimination, and restored the left NCC bridge (7/8, **Fig. 1I and J**, the signal between the arrowheads). Those pharmacological characteristics of NCC bridge formation is likely consistant with those of the [Ca^2+^]_i_ elevation (Tanaka et al., 2005). As ciliary PKD2 distribution in nodal pit appeared largely unaltered by SU5402 treatment (**Supplementary Fig. S3J and K**), the reach of PKD1L1-carrying meshwork from the NPCs toward the left ECCs, by way of the NCC bridge, may essentially trigger the Ca elevation in the left ECCs.

### Photoconverted KikGR-PKD1L1 underwent FGFR-dependent leftward shift

To test PKD1L1 protein dynamics in living embryos, we performed a photoconversion experiment in vivo. The tail buds of the knockin mouse embryos were dissected from the extraembryonic membrane, mounted on glass slides, and photobleached with a 561-nm light to eliminate the background signal in red before the experiment. Then, the nodal pit was photoconverted by a 405-nm laser light. In the consequence, red signal specifically appeared in the middle of the green area (**Fig. 2A– H; Supplementary Movie S2, 1**^**st**^ **scene**), which can better exclude the autofluorescence than the green channel. This red signal was located both on filamentous and punctate structures 5 min after the photoconversion in a control condition without inhibitor (CT; **Fig. 2G**). Interestingly, SU5402 treatment significantly decreased the number of red punctata (**Fig. 2G–I**, arrowheads; **Supplementary Movie S2, 2**^**nd**^ **scene**), suggesting that FGFR signaling may evoke a structural change on the surface of NPCs. As the exosome marker TSG-101 was also well colocalized to PKD1L1-positive punctata (**Supplementary Fig. S4A–C**), some of those punctata are likely to be exosome-like EVs. Then, we compared the global dynamics of the photoconverted signals in the absence and presence of 20 μM SU5402 in the mounting medium. 3D observation of the control embryos after 1 h of incubation significantly revealed a left-dominant appearance of linear arrays of red punctata on the endodermal cells (**Fig. 2J and L**, arrowheads; **Supplementary Movie S3**). This puncta formation was significantly eliminated by SU5402 treatment (**Fig. 2K,M,N**).

**Figure 2.**
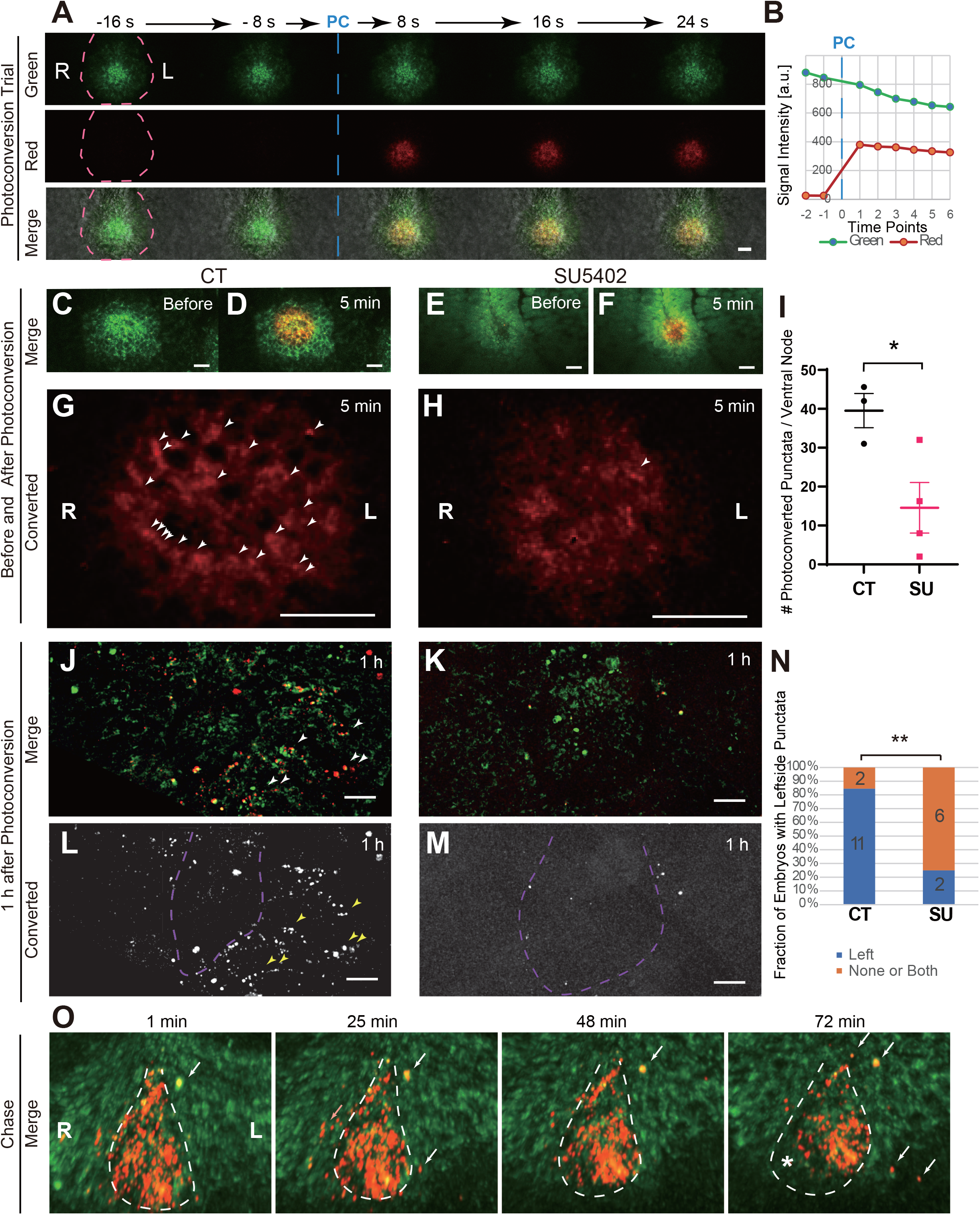
Leftward shift of KikGR-PKD1L1 protein from the nodal pit. (A and B) Double-color fluorescence micrographs during a photoconversion trial in the nodal pit of a *Pkd1l1*^*KikGR/KikGR*^ embryo at 7.5 dpc in the indicated time points (A), accompanied with statistics of the signal strengths in the NPC area (B). Scale bar, 20 μm. Dotted lines, margin of the nodal pit. Corresponding to **Supplementary Movie S2 (1**^**st**^ **scene)**. (C–I) Photoconversion in the NPCs, represented by ventral views of the embryo in low-magnified double color images (C and F) and high-magnified red color images (G and H), before (C and E) and 5 min after the photoconversion (D,F,G,H), in the absence (C,D,G) and presence of 20 μM SU5402 (E,F,H), accompanied with statistics for the number of red punctata per ventral node at 5 min after the photoconversion (I; Arrowheads in G and H). \**p* < 0.05, Welch’s *t-*test, n = 3–4. Scale bars, 20 μm. Corresponding to **Supplementary Fig. S4A–C and Movie S2 (2**^**nd**^ **scene)**. (J–N) Ventral views of embryos 1 h after photoconversion of the nodal pit, in the absence (J and L) and with 20 μM SU5402 (K and M) in double-color (J and K) and in the red channel (L and M), accompanied by statistics on the embryo fractions predominantly exhibiting red photoconverted particles (arrowheads in J and L) on the left side (N). Dotted lines, margin of the nodal pit. Scale bars, 20 μm. \*\**p* < 0.01, One-sided Fisher’s exact test with the sample numbers indicated in the graph. Corresponding to **Supplementary Movie S3**. (O) Time sequences from 4D time-lapse observation of a photoconverted *Pkd1l1*^*KikGR/KikGR*^ embryo. Dotted line, margin of the nodal pit. Arrows, punctate signals that transiently appeared on the left side of the nodal pit. Asterisk, the right half of nodal pit where the photoconverted protein had left. Corresponding to **Supplementary Movie S4**.

We further conducted long-term time-lapse experiments in an inhibitor-free condition after photoconversion in the nodal pit (**Fig. 2O, Supplementary Movie S4**). The signal within the nodal pit was reproducibly shifted to the left in 1 h, and disappeared from the right half of the nodal pit (asterisk in **Fig. 2O**). Red punctata on linear arrays transiently appeared on the left endoderm, which may be the ones expelled from the nodal pit (arrows in **Fig. 2O**). These observations collectively suggested that FGFR signaling is essential for the leftward transfer of PKD1L1 protein from the nodal pit.

### Interwoven meshwork of fibrous strands was observed in an intact preparation

To study the nodal pit in a more physiological condition, we challenged to observe intact *Pkd1l1*^*KikGR/KikGR*^ embryos directly through the extraembryonic membrane. Using differential interference contrast (DIC) microscopy, we detected the existence of layer(s) of meshwork covering the mesodermal cell surface of the nodal pit (**Fig. 3A**). Time-lapse observation of the surface of meshwork revealed EVs coming out of holes (**Fig. 3B**), which supported that the meshwork forms a “floor” in the bottom of the nodal pit. Quantitatively, large EVs (0.40 ± 0.01 μm in diameter, mean ± SEM, hereafter, n = 65) were flown out of ca. 10-μm-diameter holes of this meshwork once in ca. 3 s. The number of holes on the left half tended to be fewer than that on the right half, reflecting the presence of left NCC bridges (**Fig. 3C**). Because time-lapse analyses revealed that the EVs came out from the holes predominantly on the left half (**Fig. 3D and E; Supplementary Movies S5 and S6**), right and left holes were likely serving as the inlets and outlets of the fluid flow, respectively.

**Figure 3.**
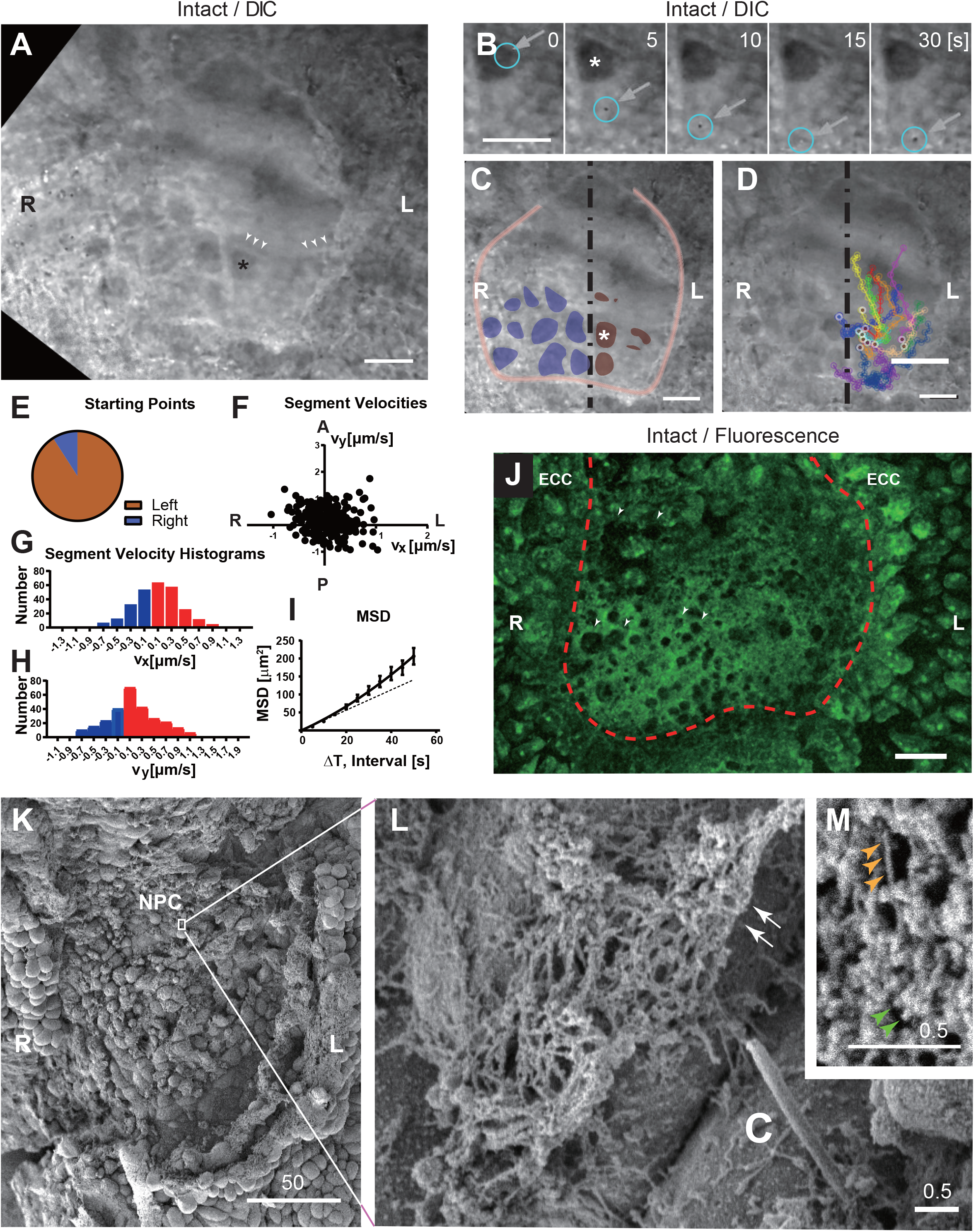
PKD1L1-containing meshwork of fibrous strands in the nodal flow. (A–I) Time lapse analysis of ventral views of an intact *Pkd1l1*^*KikGR/KikGR*^ embryo at 7.5 dpc, using a differential interference contrast (DIC) imaging on a spinning disc microscope, represented by one time frame in low magnification (A), five time frames in high magnification (B), indication of meshwork holes (blue, right; brown, left; C), particle tracking of EVs (D); accompanied by statistics of the laterality of EV-expelling holes (E), segment velocities in 2D-view (F) and in 1-D histograms along the indicated axes (G and H), and mean square displacement analysis (I). Arrowheads in (A), a ridge of meshwork. Asterisks in (A–C), the anterior left hole. Bullseye dots in (D), the first positions of the particle. Scale bars, 20 μm. Reproduced on 3 embryos. Corresponding to **Supplementary Movies S5 and S6**. (J) Fluorescent *z-*projected ventral view of an intactly-fixed *Pkd1l1*^*KikGR/KikGR*^ embryo at 7.5 dpc, Scale bar, 20 μm. Note left-dominant laterality in the ECC fluorescence level, and right-dominant laterality in the number of holes in the meshwork (arrowheads). Corresponding to **Supplementary Movie S7**. (K–M) Scanning electron micrographs of the nodal pit of a *Pkd1l1*^*KikGR/KikGR*^ embryo at 7.5 dpc that had been fixed before opening the extraembryonic membrane, at different magnifications indicated by the respective scale bars in μm. Arrows, the margin of fibrous meshwork; C, cilium; Yellow arrowheads in (M), 20-nm fibers in the core of the fibrous strand; Green arrowheads in (M), 35-nm granules associated with the fibers.

The EVs then moved toward the left wall (v_x_ = 0.08 ± 0.02 μm/s, n = 274), gradually headed to the ventral-anterior side (v_y_ = 0.14 ± 0.03 μm/s, n = 274), and then slowly returned to the right to be frequently trapped by the extraembryonic membrane (**Fig. 3F– H**). The mean square displacement curve tended to be faster than that at linear rate (at Dt = 0, the slope was 2.4 ± 0.6 μm^2^/s, n = 3, so that D = (6 ± 2) × 10^−13^ m^2^/s), consistent with the presence of active flow (**Fig. 3I**).

After fixing en bloc, the extraembryonic membrane was dissected out, and the ventral node was subjected to fluorescence microscopy. This in turn revealed the existence of fluorescent meshwork, which may be formed by PKD1L1-containing fine fibrous strands that appeared to be interwoven with one another (**Fig. 3J**). This intactly fixed preparation was further labelled for the plasma membrane lipids without any detergent treatments. Accordingly, many lamellipodia- and cytoneme-like membranous protrusions from NPCs appeared to form a part of the NCC bridge, on which PKD1L1 signal was colocalized (**Supplementary Fig. S4D–G and Movie S7**).

We further performed scanning EM of the intactly fixed fluorescent embryos (**Fig. 3K–M**). We identified a part of a huge meshwork of 10–100 nm diameter fibrous strands. They appeared to emerge from the surface of mono-ciliated nodal pit cells and from the surface of cellular protrusions (**Fig. 3K**). The size of a mesh was ca. 100–200 nm. In high-magnification view, the fibrous strands consisted of a fibrous core of 19.1 ± 0.8 nm in diameter (mean ± SEM, n = 10, **Fig. 3L**, yellow arrowheads) and the associated granules of 34.3 ± 1.8 nm in diameter (n = 10; **Fig. 3M**, green arrowheads). In whole mount immunocytochemistry, the PKD1L1-positive structure forming the NCC bridge was closely associated with but not strictly colocalizing to laminin-1-containing extracellular matrix (**Supplementary Fig. S4H–J**). It was also likely to be analogous to the recently identified de-membranous extracellular enzyme polymers termed “cytoophidium” (Liu, 2016).

### PKD1L1 was colocalized with Nodal on the left margin of the nodal pit

To scrutiny the relevance of this PKD1L1 lateralization, we double-labelled PKD1L1 and Nodal proteins in 7.5 dpc mouse embryos (**Fig. 4A–D**). The immunofluorescence of PKD1L1 protein was significantly lateralized to the left margin of nodal pit after 2–3 somite stage (9/9; **Fig. 4A**), while Nodal protein was expressed on margins on the both sides (9/9; **Fig. 4B**). Semi-superresolution microscopy further suggested colocalization of PKD1L1 and Nodal predominantly on the surface structures on the left margin (**Fig. 4C and D**).

**Figure 4.**
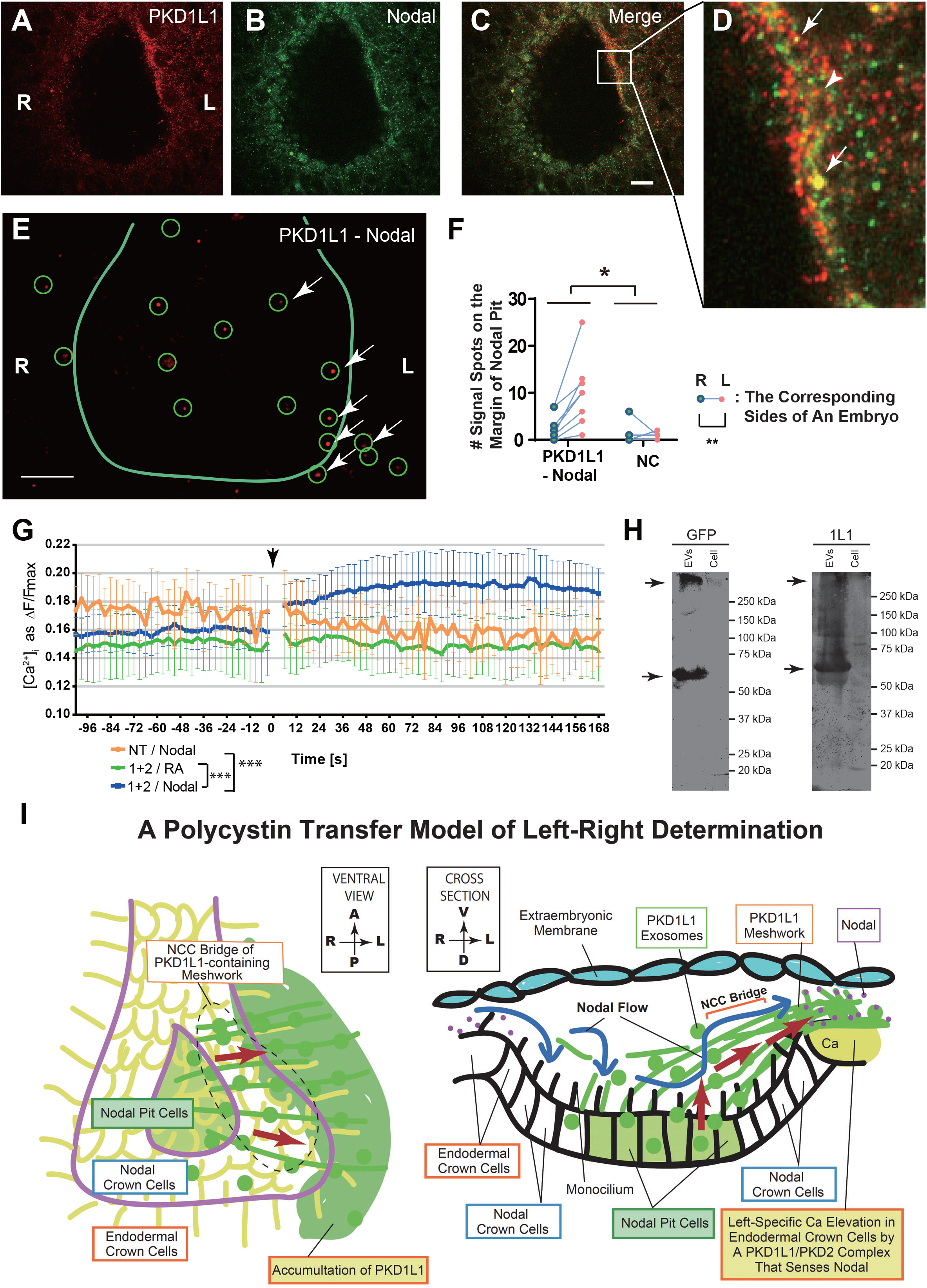
Nodal stimulates the PKD channel on the left margin of nodal pit. (A–D) An optical section of whole mount immunocytochemistry of a 4-somite stage wild type mouse embryo against PKD1L1 (red; A, C, D) and Nodal (green; B–D). Arrows, co-accumulation of PKD1L1 and Nodal on the left margin of the nodal pit. Scale bar, 10 μm. (E and F) Whole mount proximity ligation assay (PLA) against the combination of PKD1L1 and Nodal (E), accompanied with the quantification of signal spots (circled punctata with arrows) over margin of the nodal pit, on the indicated sides (F). Arrows, signal spots on the left NCCs. NC, negative control with normal IgGs. Solid line, margin of the nodal pit. Scale bar, 10 μm. \**p* < 0.05, \*\**p* < 0.01; two-way ANOVA, n = 3–8. (G) Ca transients in PKD1L1/PKD2-expressing (1+2; green and blue) or nontransfected (NT; orange) NIH3T3 cells, loaded by X-Rhod-1 AM and stimulated by 25 pM (final) recombinant mouse Nodal protein (orange and blue); or by 10^−7^ M RA (green) at time 0 (Arrow), plotted as mean ± SEM. \*\*\**p* < 0.001, two-way ANOVA during the post-stimulation period, n = 3–4. (H) Immunoblotting of the EV fraction and the whole-cell-lysate of EGFP-PKD1L1-expressing HEK293 cells against GFP and PKD1L1 (1L1). Arrows, EV condensation of co-labelled proteins in the full-length (over 300 kDa) and N-terminal half (ca. 60 kDa) forms. (I) Schematic representation of a polycystin transfer model of left-right determination. PKD1L1 is translated in NPCs and then FGFR-dependently transferred to the left, as dynamic meshwork of fibrous strands in the leftward fluid flow. On the left margin of nodal pit, condensed PKD1L1 may constitute a PKD1L1/PKD2 complex that senses Nodal to evoke long-lasting [Ca^2+^]_i_ elevation, as a readout of the nodal flow.

Then, we conducted a whole mount proximity ligation assay to reveal the association points between PKD1L1 and Nodal (**Fig. 4E**). Signal spots of association tended to appear on the left margin of the nodal pit, of which lateralization was statistically significant (**Fig. 4F**). These data suggested a left dominance of PKD1L1–Nodal interaction, consistent with the lateralized distribution of PKD1L1 protein.

### PKD1L1/PKD2-overexpressing fibroblasts became chemosensitive to Nodal protein

To test if the PKD1L1/PKD2 polycystin channel senses Nodal as a ligand, we performed calcium imaging in NIH3T3 fibroblasts. We stably transfected them with an EGFP-tagged human PKD2 expression plasmid (Doerr et al., 2016), and then transiently with a BAC vector carrying an EGFP-tagged mouse *Pkd1l1* entire gene, which significantly elevated the green fluorescence levels. Then, we measured the calcium transients using red calcium indicator, X-Rhod1 (**Fig. 4G**). Interestingly, co-expression of these polycystins enabled a long-lasting (more than 3 min) [Ca^2+^]_i_ elevation by 25 pM Nodal, but not by 10^−7^ M all-trans retinoic acid (RA). Non-transfected fibroblasts were not responding to Nodal. These data suggested that the PKD1L1 accumulation on the surface of left ECCs enhances the chemosensory polycystin channel activity there, which reacts to Nodal to evoke a long-lasting [Ca^2+^]_i_ elevation.

Finally, we analyzed the identity of N-terminal labelled PKD1L1 protein. The EV and whole cell fractions of HEK293 cells stably expressing EGFP-tagged *Pkd1l1* gene using a BAC vector were subjected to immunoblotting after stimulation by the PTEN inhibitor SF1670 (Wang et al., 2022). As a result, EGFP-PKD1L1 was significantly concentrated on the EV fraction compared with the whole cell lysates (**Fig. 4H**). Both the full-length and processed forms of PKD1L1 N-term were detected in the EV fraction, being labelled by anti-GFP and anti-PKD1L1 (N-term) antibodies at the same heights (**Fig. 4H**, arrows). These data suggested that the N-terminal tag of PKD1L1 may not be shed as a signal peptide, but chased the full-length and N-terminal polypeptides even after its extracellular release.

These data collectively suggested that the PKD1L1-containing meshwork bridges between the NPCs and left ECCs in an FGFR-dependent manner, and that PKD1L1/PKD2 polycystin serves as a receptor of Nodal (**Fig. 4I**). These working hypotheses may explain an underlying mechanism of left-dominant [Ca^2+^]_i_ elevation.

## Discussion

In this study, we have presented experimental evidence on an FGFR-dependent leftward transfer of PKD1L1 protein that may augment the Nodal chemosensory property of the polycystin channel complex on the left side of the ventral node (**Fig. 4I**), as an underlying mechanism of the left-dominant and FGFR-dependent [Ca^2+^]_i_ elevation (Tanaka *et al*., 2005). This mechanism is likely the core process of translating the laterality of nodal flow into the left-specific signal transduction in developing embryos, extending the view of “two-cilia theory” that the ciliated crown cells on the left side initiate the [Ca^2+^]_i_ elevation with the polycystin channel. The NCC bridge carrying PKD1L1 protein is lateralized to the left in an FGFR-dependent manner (**Figs. 1 and 2**), which may consist of meshwork of cellular processes, exosomes, and extracellular polymers (**Fig. 3 and supplementary Fig. S4**). PKD1L1 protein is associated with Nodal protein predominantly on the left margin of the nodal pit (**Fig. 4**). Because PKD1L1/PKD2 complex can serve as a Nodal chemosensory receptor channel in vitro (**Fig. 4G**), this left-dominant PKD1L1–Nodal interaction is a presumable process leading to the initial lateralized signal of [Ca^2+^]_i_ elevation in developing embryos.

The significance of *Pkd1l1* gene in left-right determination has been established in previous genetic studies. PKD1L1 functionally disrupting mutants, such as the medaka fish mutant strain *abc* (Kamura *et al*., 2011), the mouse mutant *rks* (Field et al., 2011), *Pkd1l1*^*-/-*^ mice (Vogel *et al*., 2010), and a human *PKD1L1-*mutation-carrying family (Le Fevre *et al*., 2020), were impaired in the proper left-right determination. Thus, its lateralization can evoke the initial [Ca^2+^]_i_ asymmetry in developing embryos. Because *Pkd1l1* mRNA itself was not expressed in a left dominant manner (**Fig. 1A**), the lateralized condensation of PKD1L1 protein may have been overlooked in previous mRNA-level studies.

Although the meshwork of PKD1L1-containing fibrous strands appeared to cover the whole area of nodal pit in the intact embryos (**Fig. 3**), the NCC bridge was robustly preserved in an extraembryonic-membrane-opened preparation (**Fig. 1**). This may represent the tight association of fibrous strands with the surface of nodal pit cells and to the left margin of the nodal pit, as speculated from the immunohistochemistry (**Fig. 1C**) and EM observations (**Fig. 3L**). These connections may be very favorable for the unidirectional transfer of the contents of the meshwork, against the eternal circulation within the nodal pit. In previous reports, cytoneme extension from *Drosophila* air sac precursor cells (Sato and Kornberg, 2002) and Shh-EV secretion from mouse limb bud mesenchyme (Wang *et al*., 2022) are highly dependent on FGFR signaling. Thus, a similar FGFR-dependent mechanisms, such as elevation of the leftward extension rate of the fibrous strands and/or their binding capacity to the left margin of the nodal pit, are expected to be involved in this NCC bridge formation. In addition, some active elimination processes of PKD1L1, such as proteolysis and/or transcytosis toward the opposite direction, are also likely to be involved in the leftward shift of PKD1L1, because the SU5402 treatment greatly enhanced the turnover of the photoconverted PKD1L1 protein (**Fig. 2M**). The elucidation of the precise function of FGFR signaling on PKD1L1 dynamics warrants further investigation.

Our experimental data on fibroblasts suggested that the PKD1L1/PKD2 complex behaves as a chemosensory channel for Nodal (**Fig. 4G**). This would be feasible because many PKD channels have been reported to serve for chemical sensing of the peptides, morphogens, and tastes (Ha *et al*., 2020; Horio *et al*., 2011; Kim *et al*., 2016). However, this argument is neutral from the long-lasting debate whether PKD channels in the ventral node can serve for mechanosensation of the fluid flow (Delling *et al*., 2016; Djenoune *et al*., 2023; Katoh *et al*., 2023; Yoshiba *et al*., 2012), because it is also likely that any mechanical stress on nodal cilia can potentiate the chemosensory properties of PKD channels. Although the *Nodal* mRNA in the ECCs has been reported to gradually lateralized to the left during the 4–5 somite stages (Mizuno *et al*., 2020), which may follow the stage of initial [Ca^2+^]_i_ elevation (**Fig. 4B**). As perturbation in calcium transients could disturb *Nodal* lateralization in 4–5 somite stage (Mizuno *et al*., 2020), the calcium laterality is likely to precede the *Nodal* laterality.

In summary, the current study will shed light on the fundamental question on how the PKD channel activity could be lateralized as a readout of the nodal flow. We presented experimental evidence for FGFR- and flow-mediated leftward shift of NCC bridge, consisting of meshwork containing a PKD channel subunit, which may then serve as a Nodal receptor on the left margin of the nodal pit. This hypothesis will nicely explain why SU5402 treatment could eliminate the [Ca^2+^]_i_ elevation, but preserving the nodal flow (Tanaka *et al*., 2005). As this study provides a good example of global dynamics of a protein that essentially serves for developmental determination, it will further stimulate the molecular and cellular biology of morphogen gradient formation in the developmental processes.

## AUTHOR CONTRIBUTION

Project administration and conceptualization by NH; Conceptualization and methodology by YT; Investigation by YT, AM; Funding acquisition by NH, YT; Original writing by YT; Review and editing by NH, AM, YT.

## ACKNOWLEDGMENTS

The NIA/NIH Mouse 15K and 7.4K cDNA Clones were provided by the RIKEN BRC through the National BioResource Project of the MEXT/AMED, Japan. *pcDNA3 GFP-h-PKD2* vector was a gift from Thomas Weimbs (Addgene plasmid #83451; http://n2t.net/addgene:83451; RRID:Addgene 83451). The authors thank Anthony A. Hyman (Max Planck Institute of Molecular Cell Biology and Genetics) for the BAC modification protocols and plasmids; Seiya Mizuno and Satoru Takahashi (Laboratory Animal Resource Center, Tsukuba University) for the knockin mouse generation; Yuuki Yamaguchi and Tomohiro Hatano (JEOL, Japan) for the SEM observation; Yoshimitsu Tsuboi (Nikon) for photoconversion; Satoru Kondo and Masumi Asahara (IRCN, The University of Tokyo) for the Nikon microscope; Shuo Wang, Reiko Takemura, Momo Morikawa, Yilong Shi, Nobuhisa Onouchi, Tsuyoshi Akamatsu, Hiromi Sato, Haruyo Fukuda, and other previous and current members of the N.H. laboratory for their technical assistance, support, and valuable discussion. This study was supported by JSPS KAKENHI grant numbers JP23000013, JP16H06372, and JP22K06246 to N.H. and by the Univ Tokyo GAP Fund (Terms #8 and #12) to Y.T.

