## Supplementary Information for "Leftward transfer of a chemosensory polycystin initiates left-dominant calcium signaling for lateralized embryonic development"

Supplementary Figures S1–S4
Supplementary Movie Legends S1–S7
Supplementary Tables S1–S3
Supplementary Methods
Supplementary References

#### SUPPLEMENTARY FIGURES

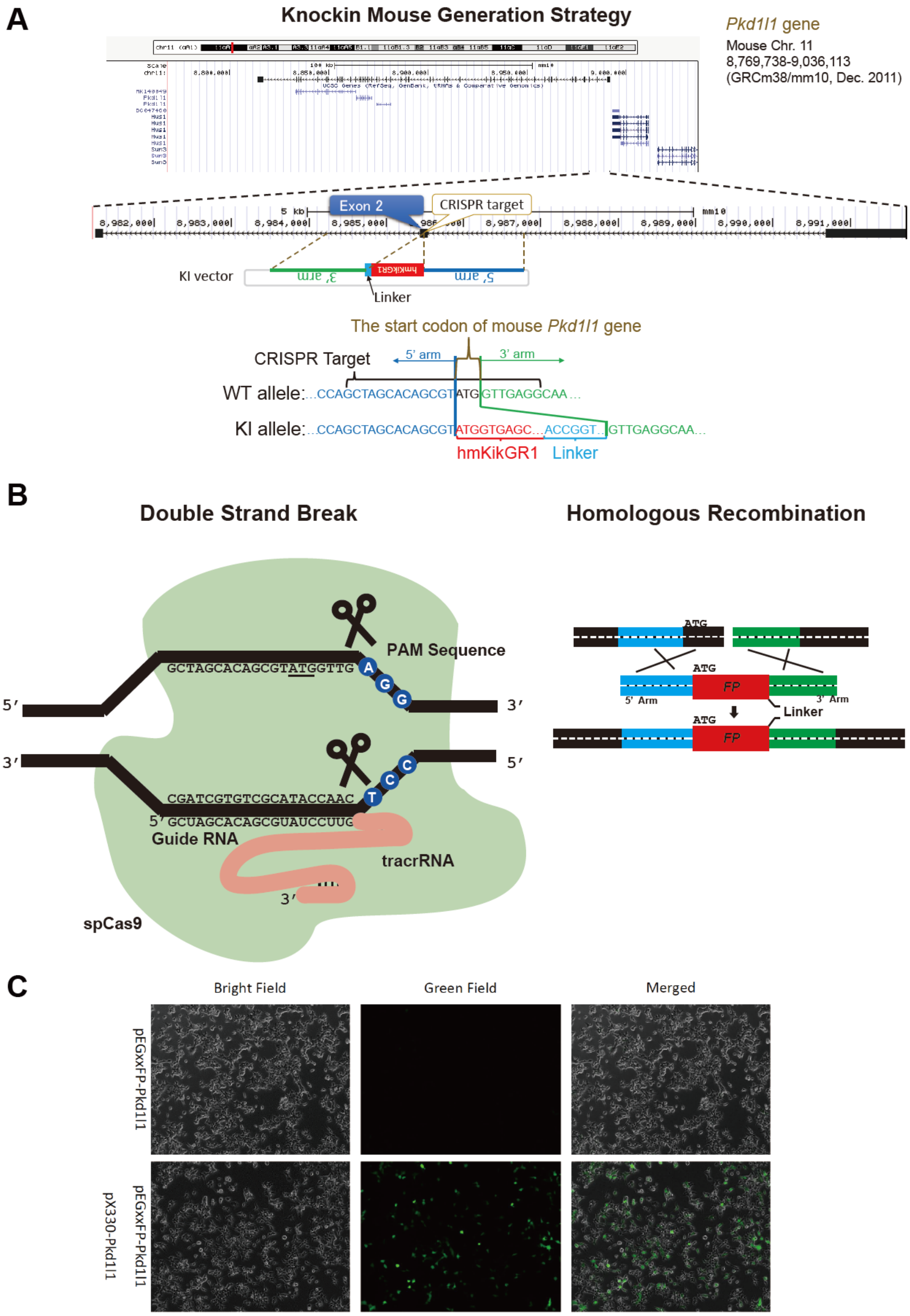

**Fig. S1. Generation of a mouse *Pkd1l1*$^{KikGR}$ allele using CRISPR/Cas9 technology, corresponding to Fig. 1E and F.**

(A) Gene editing strategy. *Pkd1l1* gene is encoded on the antisense strand of mouse Chromosome 11, and its Exon 2 contained the start codon. Using the knockin (KI) vector, a cDNA for the humanized photoconvertible fluorescent protein KikGR was knocked in just before the start codon of *Pkd1l1* gene in frame. Modified from a database search result from UCSC Genome Browser (https://genome.ucsc.edu/).

(B) Schematic representation of the double strand break and homologous recombination strategies using CRISPR/Cas9 technology.

(C) The results of a *pEGxxFP* reconstitution assay of *pX330-Pkd1l1* vector. This indicates that this vector is suitable for specifically cleaving the dsDNA in the *Pkd1l1* locus.

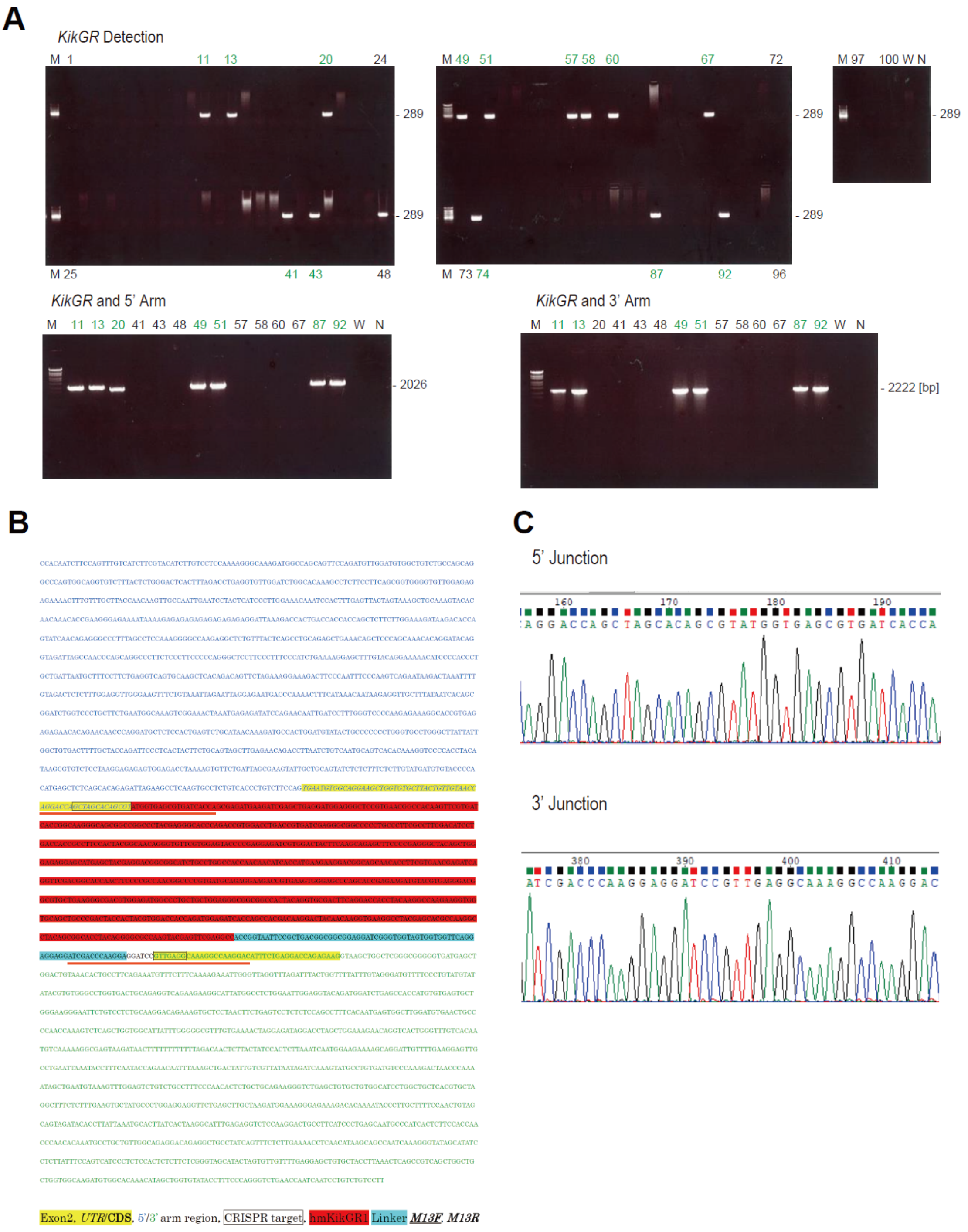


**Figure S2. Screening of *Pkd1l1*$^{KikGR}$ alleles, corresponding to Fig. 1E and F; Supplementary Tables S1–S3.**

(A) PCR genotyping of the mouse tails of the first generation after pronuclear injection of the CRISPR-Cas9 system, testing for the *KikGR* insertion and for the junctions between *hm-KikGR1* cDNA and 5'- and 3'-homologous arms, as indicated. M, molecular weight marker; W, water; N, negative control; Green numbers, the positive pups.

(B) Nucleotide sequence of the synthesized dsDNA template for homologous recombination in CRISPR/Cas9-mediated *Pkd1l1* gene editing. See the legend under the sequence for the subsequence identity. Red underlines, the sequences verified in (C).

(C) Sanger sequence electropherograms of the junctional regions.

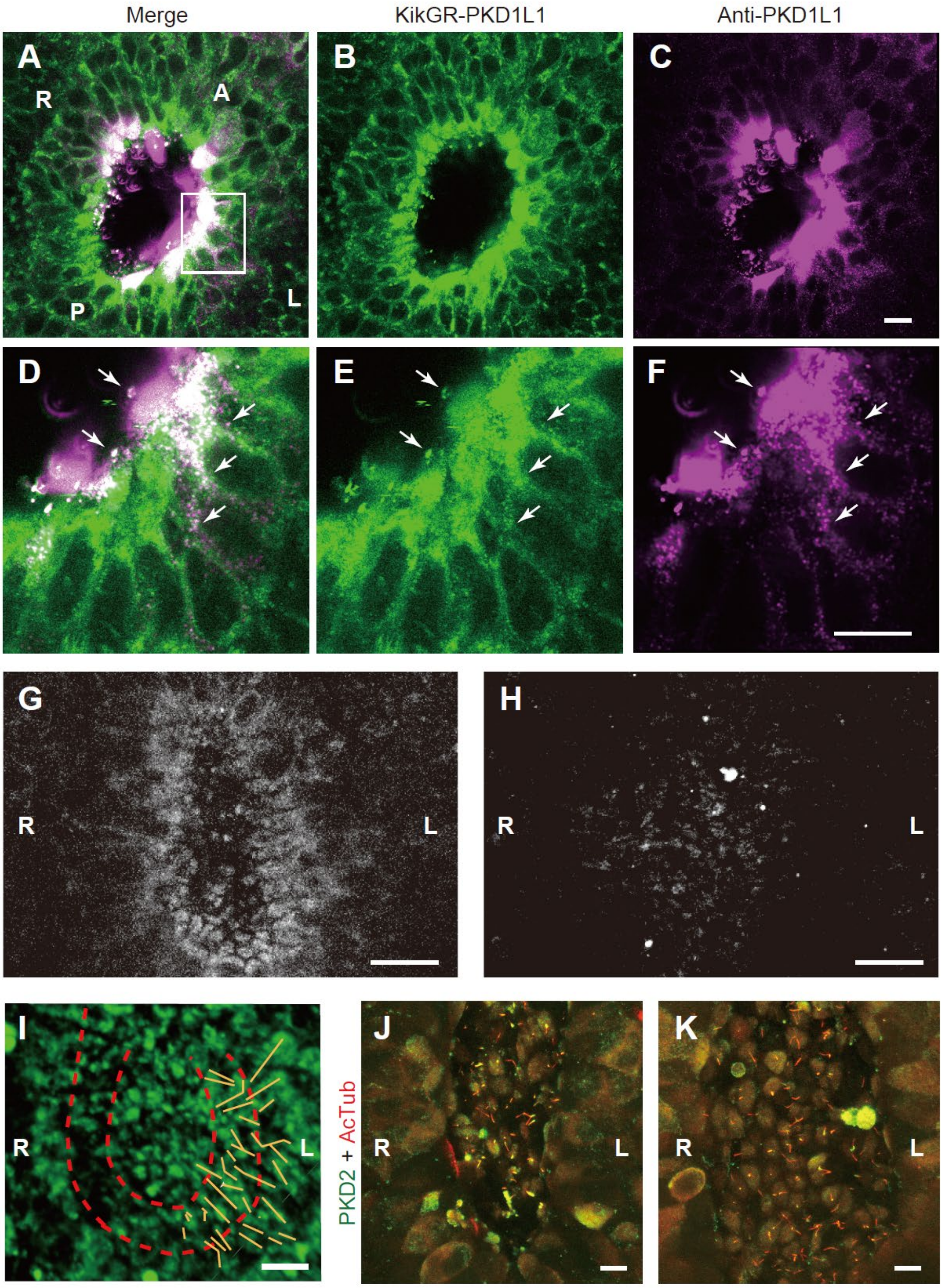


**Figure S3. Specificity of fluorescent signals in KikGR-PKD1L1 embryos, corresponding to Fig. 1G–J.**

(A–F) Correlation microscopy of the knockin tag (green, KikGR-PKD1L1; A,B,D,E) and whole mount immunohistochemistry (magenta, Anti-PKD1L1; A,C,D,F) in a single optical secretion of a *Pkd1l1*$^{KikGR/KikGR}$ embryo at low (A–C) and high (D–F) magnifications. Arrows, colocalizing spots at semi-superresolution. Scale bars, 20 µm.
(G and H) *z*-projected fluorescent micrographs in the green channel of *Pkd1l1*$^{KikGR/KikGR}$ (G) and wild-type (WT, H) embryos. Scale bars, 20 µm.
(I) Trace of strains of fluorescence signals (yellow lines) forming the left NCC bridge in the *Pkd1l1*$^{KikGR/KikGR}$ embryo in **Fig. 1G**. Dotted lines, margins of the NPC and NCC areas, respectively. Scale bar, 20 µm.
(J and K) *z*-projected whole mount immunohistochemistry of wild type mouse embryo tail buds at 7.5 dpc, treated by the carrier only (DMSO, J) and by 20 µM SU5402 (K) for 1 h, labelled against PKD2 (green) and acetylated tubulin (AcTub; red). Scale bars, 20 µm.

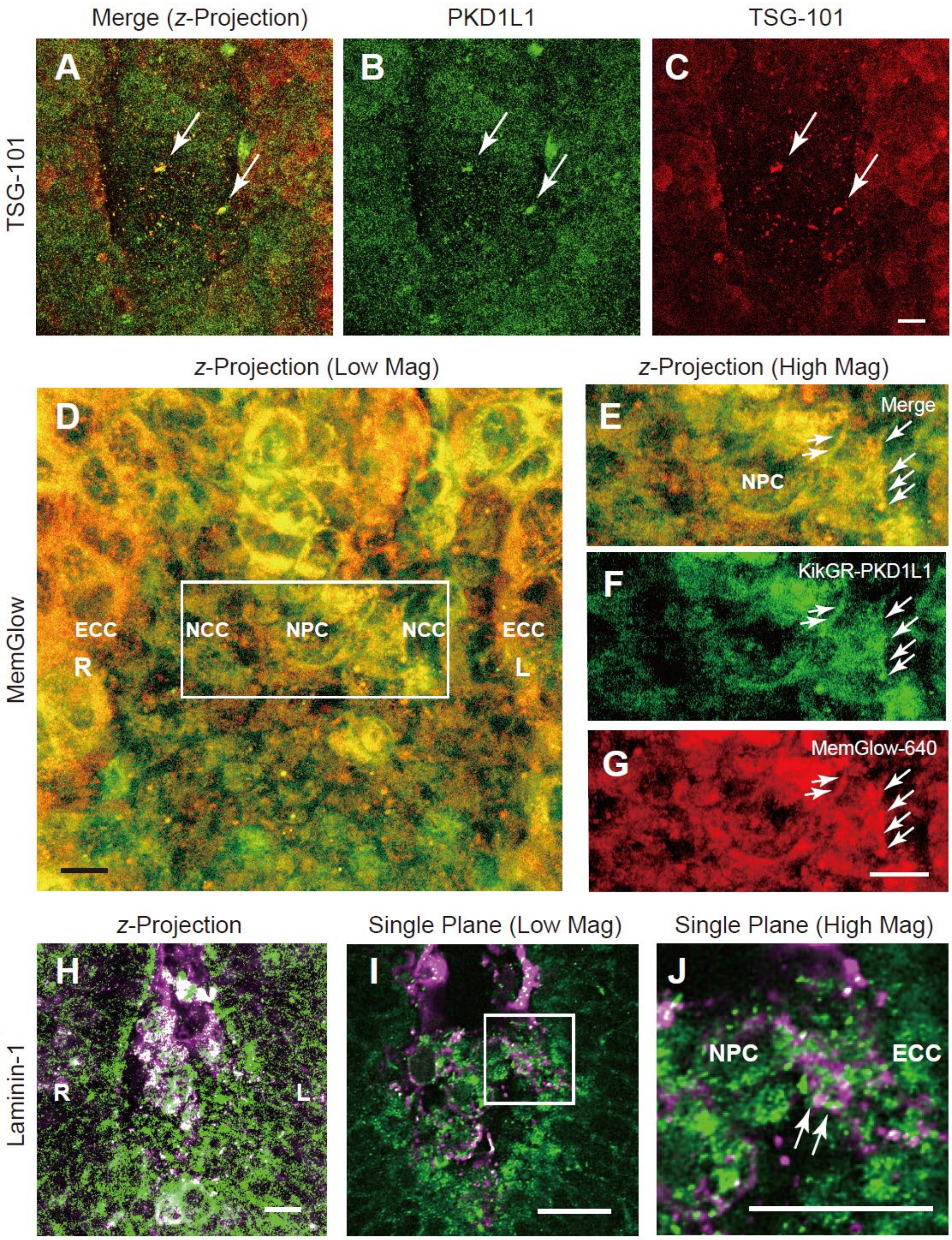


**Figure S4. Counterstaining of KikGR-PKD1L1 signals**

(A–C) *z*-projection of whole mount fluorescence microscopy of a *Pkd1l1*$^{KikGR/KikGR}$ embryo (green; A and B) at 7.5 dpc, counterstained by immunohistochemistry against the exosome marker, TSG-101 (red; A and C). Arrows, colocalization of TSG-101 and PKD1L1 to large punctata. Scale bar, 10 µm. Corresponding to **Fig. 2G.**

(D–G) *z*-projected fluorescence images of an intactly fixed *Pkd1l1*$^{KikGR/KikGR}$ embryo at 7.5 dpc, labelled in red for the cell surface with MemGlow-640 dye (red; D,E,G), overlayed with the green fluorescence image (D–F). Open box in (D) represents the magnified region in (E)–(G). Arrows, PKD1L1 signals on cytoneme-like cellular protrusions forming the left NCC bridge. Scale bars, 10 μm. Corresponding to **Fig. 3J and Supplementary Movie S7.**

(H–J) Whole-mount fluorescence microscopy of a *Pkd1l1*$^{KikGR/KikGR}$ (green) embryo, counterstained by an anti-laminin-1 antibody (magenta), in *z*-projection (H) and single plane images (I, J). Open box in (I) represents the magnified region in (J). Arrowheads, apparent colocalizations. Scale bars, 20 μm. Corresponding to **Fig. 3L**.

#### SUPPLEMENTARY MOVIE LEGENDS

**Movie S1. Laterality of KikGR-PKD1L1 signal in the nodal pit**

Rotation view of a *z*-projection of fluorescent micrographs of a $Pkd1l1^{KikGR/KikGR}$ embryo tail bud, taken by a Nikon A1R HD25 confocal microscope. Upper, anterior; Lower, posterior; Right of the movie, embryonic left side; Left of the movie, embryonic right side. Note that left dominance in NCC bridge formation (Arrow in the first scene). Corresponding to **Fig. 1G**.

**Movie S2. Photoconversion of knockin embryos**

Time-lapse images of photoconverting KikGR-PKD1L1 protein on the nodal pit cells (NPCs) of knockin embryos, where double-color fluorescence and DIC images of a control embryo are merged in the **1st scene**, and the red channel images in both conditions were presented in the **2nd scene**. Taken by a Nikon A1R HD25 confocal microscope. Upper, anterior; Lower, posterior; Right of the movie, embryonic left side; Left of the movie, embryonic right side. The whole movie of each scene consists of 12 frames of 7.8 s interval. After the first two frames, the nodal pit cell region was photoconverted from green to red according to 405-nm laser exposure. Corresponding to **Fig. 2A–H.**

**Movie S3. Appearance of KikGR-PKD1L1 protein at 1 h after photoconversion**

Rotating double-fluorescence images of photoconverted $Pkd1l1^{KikGR/KikGR}$ embryos after 1 h of treatment with 0 (Left) and 20 µM of SU5402 (Right). Taken by a Nikon A1 confocal microscope. Upper, anterior; Lower, posterior; Right of the movie, embryonic left side; Left of the movie, embryonic right side. Scale bar, 20 µm. Corresponding to **Fig. 2J–N**.

**Movie S4. Time-lapse imaging of KikGR-PKD1L1 protein after photoconversion**

*z*-projection of 4D time-lapse images of a $Pkd1l1^{KikGR/KikGR}$ embryo at 7.5 dpc, being chased for 2 h after photoconversion. Taken by a Yokogawa/ZEISS spinning disc microscope. Note that the signal within the nodal pit gradually shifts toward the left side, and that punctate signal clusters are occasionally appeared on the left side of the nodal pit. Corresponding to **Fig. 2O**.

**Movie S5. Live imaging of an intact mouse ventral node**

Time lapse image of an intact $Pkd1l1^{KikGR/KikGR}$ mouse ventral node, taken by a Yokogawa/ZEISS spinning disc microscope. Upper, anterior; Lower, posterior; Right of the movie, embryonic left side; Left of the movie, embryonic right side. Note the interwoven contour of the fibrous strands all over the ventral node, of which holes on the left side occasionally expel the EVs. On the left margin of the ventral node, cilia occasionally move through the slit between the fibrous strands. The whole movie corresponds to 2,000 s. Corresponding to **Fig. 3D**.

**Movie S6. Particle tracking in an intact mouse ventral node**
Particle tracking of the **Movie S5** by solid lines. The whole movie corresponds to 2,000 s. Corresponding to **Fig. 3D**.

**Movie S7. Localization of PKD1L1 to cellular processes on the NCC bridge**
Rotating double-fluorescence images of an intactly fixed *Pkd1l1*$^{KikGR/KikGR}$ embryo counterstained by MemGlow-640. Taken by a ZEISS LSM780-Airyscan microscope. Upper, anterior; Lower, posterior; Right of the movie, embryonic left side; Left of the movie, embryonic right side. Scale bar, 20 μm. Corresponding to **Fig. 3J** and **Supplementary Fig. S4D–G**.

### SUPPLEMENTARY TABLES

#### Supplementary Table S1. CRISPR/Cas9 Screening PCR Primer Pairs

| Screening Step | Experiment | Primers: Sequence | Primers: Primer Name | Primers: Tm (°C) | PCR Product Size (bp): WT | PCR Product Size (bp): Knock-in | PCR Product Size (bp): Random Integration: Donor Vector | PCR Product Size (bp): Random Integration: Cas9 Vector |
|---|---|---|---|---|---|---|---|---|
| 1 (Insert) | PCR | TGAACGAGATCAGGTTCGAC | KikGR detect Fw | 60 | - | 289 | 289 | - |
| | | TCGTAGGCCTTCACCTTGTT | KikGR detect Rv | 60 | | | | |
| 2 (Cas9 Tg) | PCR | AGTTCATCAAGCCCATCCTG | Cas9 detection F | 60 | - | - | - | 959 |
| | | GAAGTTTCTGTTGGCGAAGC | Cas9 detection R | 60 | | | | |
| 3 (Donor Tg) | PCR | TTGCCGGGAAGCTAGAGTAA | Amp detection-F | 57 | - | - | 560 | - |
| | | TTTGCCTTCCTGTTTTTGCT | Amp detection-R | 55 | | | | |
| 4 (3' check) | PCR | GATCGTGGACTACTTCAAGCAGAGCTTC | Pkd1l1 screening 3Fw | 68 | - | 2,222 | - | - |
| | | GCAGGTACCTTTAAAGAGCCACAGGAAA | Pkd1l1 screening 3Rv | 68 | | | | |
| | Direct Sequencing | TGAACGAGATCAGGTTCGAC | KikGR detect Fw | 50 | Joint between KikGR and ORF | | | |
| 5 (5' check) | PCR | TCGTTTTACCTGAAATCAGACTGGCTGT | Pkd1l1 screening 5Fw | 68 | - | 2,026 | - | - |
| | | AGGCCTTCACCTTGTTGTAGTCCTTGTC | Pkd1l1 screening 5Rv | 68 | | | | |
| | Direct Sequencing | AGGAGAGAGTGGAGACCTAA | Pkd1l1 screening 5 seq | 50 | Joint between UTR and KikGR | | | |

#### Supplementary Table S2. CRISPR/Cas9 Screening PCR Conditions

Step 1

| | [μl] |
|---|---|
| AmpliTaq Gold Master Mix | 10 |
| DDW | 7.8 |
| KikGR detect Fw (10 μM) | 0.6 |
| KikGR detect Rv (10 μM) | 0.6 |
| Genomic DNA (ca. 50 ng/μl) | 1 |
| Total | 20 |

| | | |
|---|---|---|
| 95°C | 10 min | |
| 95°C | 30 s | 30 Cycles |
| 60°C | 30 s | |
| 72°C | 30 s | |
| 72°C | 1.5 min | |
| 20°C | ∞ | |

Step 2

| | [μl] |
|---|---|
| AmpliTaq Gold Master Mix | 10 |
| DDW | 7.8 |
| Cas9 detection F (10 μM) | 0.6 |
| Cas9 detection R (10 μM) | 0.6 |
| Genomic DNA (ca. 50 ng/μl) | 1 |
| Total | 20 |

| | | |
|---|---|---|
| 95°C | 10 min | |
| 95°C | 30 s | 35 Cycles |
| 60°C | 30 s | |
| 72°C | 1 min | |
| 72°C | 3 min | |
| 20°C | ∞ | |

Step 3

| | [μl] |
|---|---|
| AmpliTaq Gold Master Mix | 10 |
| DDW | 7.8 |
| Amp detection-F (10 μM) | 0.6 |
| Amp detection-R (10 μM) | 0.6 |
| Genomic DNA (ca. 50 ng/μl) | 1 |
| Total | 20 |

| | | |
|---|---|---|
| 95°C | 10 min | |
| 95°C | 30 s | 30 Cycles |
| 58°C | 30 s | |
| 72°C | 45 s | |
| 72°C | 2 min | |
| 20°C | ∞ | |

Step 4

| | [μl] |
|---|---|
| DDW | 18.6 |
| Primestar GXL | 0.6 |
| Buffer | 6 |
| dNTP | 2.4 |
| Pkd1l1 screening 3Fw (10 μM) | 0.7 |
| Pkd1l1 screening 3Rv (10 μM) | 0.7 |
| Genomic DNA (ca. 50 ng/μl) | 1 |
| Total | 30 |

| | | |
|---|---|---|
| 98°C | 2 min | |
| 98°C | 10 s | 30 Cycles |
| 68°C | 2.5 min | |
| 68°C | 7.5 min | |
| 4°C | ∞ | |

Step 5

| | [μl] |
|---|---|
| DDW | 18.6 |
| Primestar GXL | 0.6 |
| Buffer | 6 |
| dNTP | 2.4 |
| Pkd1l1 screening 5Fw (10 μM) | 0.7 |
| Pkd1l1 screening 5Rv (10 μM) | 0.7 |
| Genomic DNA (ca. 50 ng/μl) | 1 |
| Total | 30 |

| | | |
|---|---|---|
| 98°C | 2 min | |
| 98°C | 10 s | 30 Cycles |
| 68°C | 2.5 min | |
| 68°C | 7.5 min | |
| 4°C | ∞ | |

### Table S3. F1 Mouse Genotyping

| Pkd1l1 KikGR KI # | Sex | Knock-in detection | | | | | Random Tg Detection | |
|---|---|---|---|---|---|---|---|---|
| | | KikGR detection PCR | 5' PCR (5'arm out to insert) | 3' PCR (insert to 3'arm out) | Joint Sequence | | Cas9 dection PCR | Amp$^R$ detection PCR |
| | | | | | 5'UTR to KIKGR | KIKGR to Pkd1l1 ORF | | |
| 1 | ♂ | × | Not Examined | Not Examined | Not Examined | Not Examined | Not Examined | Not Examined |
| 2 | ♂ | × | Not Examined | Not Examined | Not Examined | Not Examined | Not Examined | Not Examined |
| 3 | ♂ | × | Not Examined | Not Examined | Not Examined | Not Examined | Not Examined | Not Examined |
| 4 | ♂ | × | Not Examined | Not Examined | Not Examined | Not Examined | Not Examined | Not Examined |
| 5 | ♂ | × | Not Examined | Not Examined | Not Examined | Not Examined | Not Examined | Not Examined |
| 6 | ♂ | × | Not Examined | Not Examined | Not Examined | Not Examined | Not Examined | Not Examined |
| 7 | ♂ | × | Not Examined | Not Examined | Not Examined | Not Examined | Not Examined | Not Examined |
| 8 | ♂ | × | Not Examined | Not Examined | Not Examined | Not Examined | Not Examined | Not Examined |
| 9 | ♂ | × | Not Examined | Not Examined | Not Examined | Not Examined | Not Examined | Not Examined |
| 10 | ♂ | × | Not Examined | Not Examined | Not Examined | Not Examined | Not Examined | Not Examined |
| 11 | ♂ | ✔ | ✔ | ✔ | ✔ | ✔ | Negative | Negative |
| 12 | ♂ | × | Not Examined | Not Examined | Not Examined | Not Examined | Not Examined | Not Examined |
| 13 | ♂ | ✔ | ✔ | ✔ | ✔ | ✔ | Negative | Negative |
| 14 | ♂ | × | Not Examined | Not Examined | Not Examined | Not Examined | Not Examined | Not Examined |
| 15 | ♂ | × | Not Examined | Not Examined | Not Examined | Not Examined | Not Examined | Not Examined |
| 16 | ♂ | × | Not Examined | Not Examined | Not Examined | Not Examined | Not Examined | Not Examined |
| 17 | ♂ | × | Not Examined | Not Examined | Not Examined | Not Examined | Not Examined | Not Examined |
| 18 | ♂ | × | Not Examined | Not Examined | Not Examined | Not Examined | Not Examined | Not Examined |
| 19 | ♂ | × | Not Examined | Not Examined | Not Examined | Not Examined | Not Examined | Not Examined |
| 20 | ♂ | ✔ | × | × | Not Examined | Not Examined | Negative | Positive |
| 21 | ♂ | × | Not Examined | Not Examined | Not Examined | Not Examined | Not Examined | Not Examined |
| 22 | ♂ | × | Not Examined | Not Examined | Not Examined | Not Examined | Not Examined | Not Examined |
| 23 | ♂ | × | Not Examined | Not Examined | Not Examined | Not Examined | Not Examined | Not Examined |
| 24 | ♂ | × | Not Examined | Not Examined | Not Examined | Not Examined | Not Examined | Not Examined |
| 25 | ♂ | × | Not Examined | Not Examined | Not Examined | Not Examined | Not Examined | Not Examined |
| 26 | ♂ | × | Not Examined | Not Examined | Not Examined | Not Examined | Not Examined | Not Examined |
| 27 | ♂ | × | Not Examined | Not Examined | Not Examined | Not Examined | Not Examined | Not Examined |
| 28 | ♂ | × | Not Examined | Not Examined | Not Examined | Not Examined | Not Examined | Not Examined |
| 29 | ♂ | × | Not Examined | Not Examined | Not Examined | Not Examined | Not Examined | Not Examined |
| 30 | ♂ | × | Not Examined | Not Examined | Not Examined | Not Examined | Not Examined | Not Examined |
| 31 | ♂ | × | Not Examined | Not Examined | Not Examined | Not Examined | Not Examined | Not Examined |
| 32 | ♂ | × | Not Examined | Not Examined | Not Examined | Not Examined | Not Examined | Not Examined |
| 33 | ♂ | × | Not Examined | Not Examined | Not Examined | Not Examined | Not Examined | Not Examined |
| 34 | ♂ | × | Not Examined | Not Examined | Not Examined | Not Examined | Not Examined | Not Examined |
| 35 | ♂ | × | Not Examined | Not Examined | Not Examined | Not Examined | Not Examined | Not Examined |
| 36 | ♂ | × | Not Examined | Not Examined | Not Examined | Not Examined | Not Examined | Not Examined |
| 37 | ♂ | × | Not Examined | Not Examined | Not Examined | Not Examined | Not Examined | Not Examined |
| 38 | ♂ | × | Not Examined | Not Examined | Not Examined | Not Examined | Not Examined | Not Examined |
| 39 | ♂ | × | Not Examined | Not Examined | Not Examined | Not Examined | Not Examined | Not Examined |
| 40 | ♂ | × | Not Examined | Not Examined | Not Examined | Not Examined | Not Examined | Not Examined |
| 41 | ♂ | ✔ | × | × | Not Examined | Not Examined | Negative | Negative |
| 42 | ♂ | × | Not Examined | Not Examined | Not Examined | Not Examined | Not Examined | Not Examined |
| 43 | ♂ | ✔ | × | × | Not Examined | Not Examined | Negative | Negative |
| 44 | ♀ | × | Not Examined | Not Examined | Not Examined | Not Examined | Not Examined | Not Examined |
| 45 | ♀ | × | Not Examined | Not Examined | Not Examined | Not Examined | Not Examined | Not Examined |
| 46 | ♀ | × | Not Examined | Not Examined | Not Examined | Not Examined | Not Examined | Not Examined |
| 47 | ♀ | × | Not Examined | Not Examined | Not Examined | Not Examined | Not Examined | Not Examined |
| 48 | ♀ | ✔ | × | × | Not Examined | Not Examined | Negative | Negative |
| 49 | ♀ | ✔ | ✔ | ✔ | ✔ | ✔ | Negative | Positive |
| 50 | ♀ | × | Not Examined | Not Examined | Not Examined | Not Examined | Not Examined | Not Examined |
| 51 | ♀ | ✔ | ✔ | ✔ | ✔ | ✔ | Negative | Positive |
| 52 | ♀ | × | Not Examined | Not Examined | Not Examined | Not Examined | Not Examined | Not Examined |
| 53 | ♀ | × | Not Examined | Not Examined | Not Examined | Not Examined | Not Examined | Not Examined |
| 54 | ♀ | × | Not Examined | Not Examined | Not Examined | Not Examined | Not Examined | Not Examined |
| 55 | ♀ | × | Not Examined | Not Examined | Not Examined | Not Examined | Not Examined | Not Examined |
| 56 | ♀ | × | Not Examined | Not Examined | Not Examined | Not Examined | Not Examined | Not Examined |
| 57 | ♀ | ✔ | × | × | Not Examined | Not Examined | Negative | Negative |
| 58 | ♀ | ✔ | × | × | Not Examined | Not Examined | Negative | Negative |
| 59 | ♀ | × | Not Examined | Not Examined | Not Examined | Not Examined | Not Examined | Not Examined |
| 60 | ♀ | ✔ | × | × | Not Examined | Not Examined | Negative | Negative |
| 61 | ♀ | × | Not Examined | Not Examined | Not Examined | Not Examined | Not Examined | Not Examined |
| 62 | ♀ | × | Not Examined | Not Examined | Not Examined | Not Examined | Not Examined | Not Examined |
| 63 | ♀ | × | Not Examined | Not Examined | Not Examined | Not Examined | Not Examined | Not Examined |
| 64 | ♀ | × | Not Examined | Not Examined | Not Examined | Not Examined | Not Examined | Not Examined |
| 65 | ♀ | × | Not Examined | Not Examined | Not Examined | Not Examined | Not Examined | Not Examined |
| 66 | ♀ | × | Not Examined | Not Examined | Not Examined | Not Examined | Not Examined | Not Examined |
| 67 | ♀ | ✔ | × | × | Not Examined | Not Examined | Negative | Not Examined |
| 68 | ♀ | × | Not Examined | Not Examined | Not Examined | Not Examined | Not Examined | Not Examined |
| 69 | ♀ | × | Not Examined | Not Examined | Not Examined | Not Examined | Not Examined | Not Examined |
| 70 | ♀ | × | Not Examined | Not Examined | Not Examined | Not Examined | Not Examined | Not Examined |
| 71 | ♀ | × | Not Examined | Not Examined | Not Examined | Not Examined | Not Examined | Not Examined |
| 72 | ♀ | × | Not Examined | Not Examined | Not Examined | Not Examined | Not Examined | Not Examined |
| 73 | ♀ | × | Not Examined | Not Examined | Not Examined | Not Examined | Not Examined | Not Examined |
| 74 | ♀ | ✔ | Not Examined | Not Examined | Not Examined | Not Examined | Positive | Not Examined |
| 75 | ♀ | × | Not Examined | Not Examined | Not Examined | Not Examined | Not Examined | Not Examined |
| 76 | ♀ | × | Not Examined | Not Examined | Not Examined | Not Examined | Not Examined | Not Examined |
| 77 | ♀ | × | Not Examined | Not Examined | Not Examined | Not Examined | Not Examined | Not Examined |
| 78 | ♀ | × | Not Examined | Not Examined | Not Examined | Not Examined | Not Examined | Not Examined |
| 79 | ♀ | × | Not Examined | Not Examined | Not Examined | Not Examined | Not Examined | Not Examined |
| 80 | ♀ | × | Not Examined | Not Examined | Not Examined | Not Examined | Not Examined | Not Examined |
| 81 | ♀ | × | Not Examined | Not Examined | Not Examined | Not Examined | Not Examined | Not Examined |
| 82 | ♀ | × | Not Examined | Not Examined | Not Examined | Not Examined | Not Examined | Not Examined |
| 83 | ♀ | × | Not Examined | Not Examined | Not Examined | Not Examined | Not Examined | Not Examined |
| 84 | ♀ | × | Not Examined | Not Examined | Not Examined | Not Examined | Not Examined | Not Examined |
| 85 | ♀ | × | Not Examined | Not Examined | Not Examined | Not Examined | Not Examined | Not Examined |
| 86 | ♀ | × | Not Examined | Not Examined | Not Examined | Not Examined | Not Examined | Not Examined |
| 87 | ♀ | ✔ | ✔ | ✔ | ✔ | ✔ | Negative | Positive |
| 88 | ♀ | × | Not Examined | Not Examined | Not Examined | Not Examined | Not Examined | Not Examined |
| 89 | ♀ | × | Not Examined | Not Examined | Not Examined | Not Examined | Not Examined | Not Examined |
| 90 | ♀ | × | Not Examined | Not Examined | Not Examined | Not Examined | Not Examined | Not Examined |
| 91 | ♀ | × | Not Examined | Not Examined | Not Examined | Not Examined | Not Examined | Not Examined |
| 92 | ♀ | ✔ | ✔ | ✔ | ✔ | ✔ | Negative | Negative |
| 93 | ♀ | × | Not Examined | Not Examined | Not Examined | Not Examined | Not Examined | Not Examined |
| 94 | ♀ | × | Not Examined | Not Examined | Not Examined | Not Examined | Not Examined | Not Examined |
| 95 | ♀ | × | Not Examined | Not Examined | Not Examined | Not Examined | Not Examined | Not Examined |
| 96 | ♀ | × | Not Examined | Not Examined | Not Examined | Not Examined | Not Examined | Not Examined |
| 97 | ♀ | × | Not Examined | Not Examined | Not Examined | Not Examined | Not Examined | Not Examined |
| 98 | ♀ | × | Not Examined | Not Examined | Not Examined | Not Examined | Not Examined | Not Examined |
| 99 | ♀ | × | Not Examined | Not Examined | Not Examined | Not Examined | Not Examined | Not Examined |
| 100 | ♀ | × | Not Examined | Not Examined | Not Examined | Not Examined | Not Examined | Not Examined |

#### SUPPLEMENTARY METHODS

**Mouse knockin.** *Pkd1l1*$^{KikGR/KikGR}$ mice were produced at Transborder Medical Research Center, University of Tsukuba, basically using previously described methods of CRISPR-Cas9 technology (Mashiko et al., 2013; Nakagawa et al., 2016) as depicted in **Supplementary Fig. 1A and B**. The first ATG in Exon 2 of the mouse *Pkd1l1* gene on Chromosome 11 was substituted by a humanized monomeric *KikGR1* + Linker + *BamHI* sequence. To achieve a double-strand break, the CRISPR target 5'-GCTAGCACAGCGTATGGTTG**AGG**-3' (underline, the first ATG; bold, the PAM sequence; chr11:8985496–8985474) was inserted into the Cas9-expressing *pX330* vector (Addgene plasmid #42230). The double-strand-break-mediated homology-dependent repair efficiency of this *pX330-Pkd1l1* vector was verified by a *pEGxxFP* reconstitution assay (**Supplementary Fig. S1C**) (Mashiko *et al.*, 2013). A homologous recombination template was generated by synthesizing a 3,311-bp-long dsDNA of *hmKikGR1* + Linker + *BamHI* sequence flanked by 5'- and 3'-homologous arms on a *pUC57-Amp* vector (Genewiz). 10 ng/ml of recombination template plasmid and 5 ng/ml of *pX330-PKd1l1* vector were simultaneously injected into the pronuclei of one-cell-stage C57BL/6J mouse embryos, which were transferred into pseudopregnant ICR mouse oviducts. Three out of one hundred pups were identified to be positive for homologous recombination without random integrations according to genomic PCR and Sanger sequencing that was 100% match to the expected sequence (**Supplementary Fig. S1B–D** and **Supplementary Tables S1–S3**). Their offspring were maintained in a C57BL/6J background in specific pathogen-free environment under a 14/10-hr light/dark cycle, in accordance with the institutional guidelines and approval of University of Tsukuba and The University of Tokyo. Homozygote embryos at 7.5 dpc were routinely obtained by intercrossing the homozygote mouse pairs.

**Whole mount embryo culture**. Whole mount embryo culture was performed basically as previously described (Sturm and Tam, 1993). Briefly, mouse embryos were released from decidua in PB1/10% FCS medium and the extraembryonic membrane was opened. They were then incubated in DR75 medium at 37°C in 5% $CO_2$ atmosphere.

**EGFP tagging of mouse *Pkd1l1* gene in BAC.** Bacterial artificial chromosome (BAC) clone RPC1-23-39H1 (Chr 11, 8,752,177–8,987,803 nt) in *E.coli* was purchased from DNAFORM, on the reverse strand of which *Pkd1l1* was encoded on 8,973,266–8,826,708 nt according to the BACfinder website (www.mitocheck.org/cgi-bin/BACfinder). It was transformed with *pRedET* (GeneBridge) by electroporation, then with a homologous recombination template being amplified from *R6Kamp-hNGFP* cassette (kindly provided by Dr. Anthony A. Hyman) with the following primers *NFLAPfwd*: 5'-CAGGAAGCTGGTGTGCTTACTGTTGTAACCAGGACCAGCTAGCACAGCGTAT GGTGTCCAAGGGCGAGGAACTG-3' and *NFLAPrev*: 5'-GCTTACCTTCTCTGGTCCTCAGAAATGTCCTTGGCCTTTGCCTCAACCATGGC CCTGGGCAGGTCGTCGGTCAG-3' using KOD-Plus-Neo DNA polymerase (Toyobo). The homologous recombinant clones were screened by PCR using the following primers: F1: 5'-GTTCACATCCAAGCCACTCATTG-3' and B1: 5'-GCCGCCAAGTTCAAAGAAACAG-3' and clone #4-1 was turned out to be positive. It was subjected to large scale DNA preparation using a NucleoBond Xtra BAC kit (Takara Bio) and transfected into fibroblasts using Lipofectamine LTX Plus reagent (Thermo Fisher) according to the manufacturers' protocols.

**Whole mount in situ hybridization**. Whole mount in situ hybridization was performed basically as previously described (Nonaka *et al.*, 1998), with a 10 µg/ml Protease K treatment for 5 min. A fragment of mouse *Pkd1l1* cDNA (4,735–8,928 bp) (Tamplin *et al.*, 2011) subcloned in *pSPORT1* vector was purchased from RIKEN Bioresource Research Center (NIA/NIH Mouse cDNA Clone #H3025A11), of which insert was amplified by PCR using the M13 universal primers. Using the PCR product as a template, a digoxigenin (DIG)-labelled riboprobe was transcribed with SP6 RNA polymerase using a DIG RNA Labelling Mix (Sigma-Aldrich) and purified using a Chroma Spin TM-1000 column (Takara Bio), to be subjected to hybridization.

**Antibodies and fluorescent reagents.** An anti-PKD1L1 polyclonal antibody (#OSP00015W; RRID:AB_2163361; 1:200) was purchased from Osenses Pty Ltd; an anti-Nodal polyclonal antibody (#AF1315; RRID:AB_2151531) and anti-laminin-1 monoclonal antibody (#MAB2549; RRID:AB_2133911) were from R & D Systems; an anti-TSG-101 monoclonal antibody (#MA1-23296, RRID:AB_2208088), normal rabbit IgG (#ICN55944; RRID:AB_2334717), and normal mouse IgG (#02-6502; RRID:AB_2532951) were from Thermo Fisher Scientific; a rabbit polyclonal anti-GFP antibody (#598; RRID:AB_591816) was from MBL International; and MemGlow-640 Fluorogenic Plasma Membrane Probe (#MG04-10) was from Cytoskeleton. Alexa-Fluor-labelled secondary antibodies were purchased from Thermo Fisher Scientific and used at 1:500.

**Whole mount fluorescent microscopy**. Whole mount immunohistochemistry was performed basically as previously described (Tanaka *et al.*, 2005). For conventional fixation, the tail bud of an ICR mouse embryo at 7.5 dpc was dissected in 10% FBS/PB1, washed in PBS, fixed in 4% paraformaldehyde/PBS, permeabilized with 0.1% Triton-X 100 in PBS at room temperature (RT) for 5 min, treated with the primary antibody at 4ºC overnight and the secondary antibody at RT for 1 h in Can Get Signal solution (TOYOBO) with brief washing steps in PBS, and mounted with a coverslip and 0.2-mm-thick silicon spacer. They were subjected to confocal microscopy with an LSM780 Airyscan (ZEISS), a spinning disc confocal microscopy (ZEISS-Yokogawa), equipped with a 40×/1.1 C-Apochromat long-working-distance water immersion objective lens (ZEISS) as previously described (Tanaka et al., 2016), or Nikon A1 confocal microscopy at IRCN, Univ Tokyo. For observing the linear arrays of fluorescent signals in the nodal pit, the knockin embryos were dissected out from the decidua carefully avoiding disruption of the extraembryonic membrane, then soaked in the half Kalnovsky fixative or 2% paraformaldehyde/0.1% glutaraldehyde/PBS at 37ºC for 10 min and 4ºC for overnight, soaked in 0.1–0.5% paraformaldehyde in PBS for overnight, dissected out the extraembryonic membrane in 10% FBS/PB1 by fine forceps, and postfixed to be subjected to microscopy. Either the *z*-projection image or the single optical section image was presented as described in the figure legends.

**DIC observation in vivo.** For observing the ventral node in an intact preparation, the embryos at 7.5 dpc were dissected out of the decidua *en bloc* in PB1–10% FCS. They were mounted on a chambered coverslip (ibidi, #80486) and directly observed with a ZEISS/Yokogawa spinning disc microscope equipped with a 40×/1.1 C-Apochromat long-working-distance water immersion objective lens (ZEISS) and a differential interference contrast (DIC) system in a time-lapse manner at a frame rate of 0.2–25 frame per second. The obtained images were manually tracked using the MTrackJ (Meijering et al., 2012) plugin on ImageJ/Fiji software (Schneider et al., 2012) to present a particle-tracking movie and to measure the segment velocities that were subjected to the scattered plot, histogram, and mean square displacement analyses using the Prism 9.3.1 software (GraphPad Software).

**Surface staining.** For staining the surface, fixed embryos were soaked in freshly diluted MemGlow-640 solution in PBS (1:1,000) for 10 min according to the manufacturer's protocol, and subjected to a ZEISS LSM780-Airyscan microscope.

**Whole mount proximity ligation assay**. For whole mount proximity ligation assay, the mouse embryo tail bud was dissected, fixed, permeabilized, and treated with the primary antibodies as described in the above. They were then subjected to proximity ligation assay using a Duolink kit (Sigma Aldrich) according to the manufacturer's protocol, mounted on glass slides, and observed with confocal microscopy. The number of punctate signals on the margin of nodal pit was counted for statistical analyses.

**Scanning Electronmicroscopy (SEM).** SEM preparation was performed basically as previously described (Nonaka *et al.*, 1998), but the chemical fixation with Half-Karnovsky fixative was performed before opening the extraembryonic membrane. After the fluorescence was verified using confocal laser scanning microscopy, dehydrated and isoamyl-acetate-soaked samples were subjected to critical point drying with liquid $CO_2$, coated by an Osmium coater (JEOL), and observed with a JSM-IT800 scanning EM (JEOL).

**In vitro Ca imaging.** For measuring Ca transient, NIH3T3 cells were stably transfected with a *pcDNA3 GFP-h-PKD2* vector (Doerr et al., 2016)(Addgene #83451) with Lipofectamine LTX Plus (Thermo Fisher) and selected by 200 μg/ml hygromycin B (Invitrogen), whose GFP fluorescence was verified by fluorescence microscopy. Then, those PKD2-expressing cells were transfected with the *EGFP-Pkd1l1* knockin BAC clone #4-1 in a transient manner, which was identified by significant enhancement of the EGFP fluorescence. The cells were loaded with the Ca indicator X-Rhod-1 AM (Thermo Fisher, # X14210) at 5 μg/ml for 15 min and subjected to a ZEISS LSM780 confocal laser-scanning microscope equipped with a stage-top $CO_2$ incubator. The green fluorescence excited by 488-nm laser was detected by the lambda mode, and well-differentiated and doubly transfected cells with the mean fluorescence level exceeding 11, larger than 400 $\mu m^2$, and with a stable baseline with the 568-nm excitation were selected. After 50 scans every 5 s, recombinant mouse Nodal protein (R&D systems, #1315-ND) was supplemented at 25 pM, and chased for another 50 frames. The mean $\Delta F/F_{max}$ values were plotted after removing outliers at more than mean + 2 S.D. and compared with those of nontransfected controls, according to the previously described methods (Tanaka *et al.*, 2016).

**Photoconversion.** *Pkd1l1*$^{KikGR/KikGR}$ mouse embryos at 7.5 dpc were recovered from decidua, and cultured in DR75 medium (phenol-red-free) for 10 min to several hours after the extraembryonic membrane was opened. Their tail buds were then dissected in PB1/FBS and mounted on glass slides as previously described (Tanaka *et al.*, 2005). They were then subjected to a Nikon A1R HD25 confocal laser-scanning microscope equipped with a 60×/1.2 Plan-Apo VC oil-immersion objective lens. Their NPCs were three times exposed to 405-nm laser light at 20%, scan speed 1 (pixel dwell 1.9), averaged for 8 times, in a region of interest from 512 × 512 pixels, to ensure that a significantly red-converted region appeared (**Fig. 2A–F**). They were then incubated in a dark box at room temperature for more than 60 min, to be subjected to 3D or 4D observation using a spinning disc confocal microscope that is an Axio Observer inverted microscope (ZEISS) equipped with a long-working-distance water-immersion C-Apochromat lens (40×, NA 1.1) and a CSU-W1 confocal scanner unit (Yokogawa). 3D reconstruction of those obtained *z*-stacks were performed using ImageJ 1.52n (https://imagej.nih.gov/ij/docs/index.html) or NIS-Elements AR 5.11.0 (Nikon) software with a deconvolution mode. For long-term observations, we have used a resonance mode of a Nikon A1 confocal microscope at IRCN, The University of Tokyo, with 40×/1.25 water-immersion Apochromat objective lens at 512 × 512 pixels at the speed of 7.5 frame/s averaged for 32 times. After the 4D imaging at 3-μm intervals for 2 h, the images were deconvoluted and z-projected, and subjected for analyses.

**In vitro pharmacology.** The SU5402 and/or Shh-N treatments of mouse embryos was performed as described previously (Tanaka *et al.*, 2005). Whole mount embryos were dissected out of the decidua with the extraembryonic membrane opened and cultured in DR75 medium with or without 20 μM SU5402 (Sigma Aldrich) and/or 2 μg/ml recombinant mouse Shh-N peptide (#461-SH-025, R&D Systems) for 1 h at 37ºC with 5% $CO_2$. The tail buds were then dissected from the embryos, mounted, and subjected to 3D or 4D observation using a ZEISS/Yokogawa spinning disc microscope, a ZEISS LSM780-Airyscan confocal laser scanning microscope, and a Nikon A1R HD25 confocal laser scanning microscope.

**Immunoblotting of the EV fraction.** HEK293 cells were stably transfected with *EGFP-Pkd1l1* gene-containing BAC clone with 200 μg/ml G418 (Thermo Fisher). The culture was stimulated overnight by the PTEN inhibitor SF1670 as described previously (Wang *et al.*, 2022), and the conditioned medium was subjected to EV precipitation using Total Exosome Isolation Reagent (#4478359, Thermo Fisher) following the manufacturer's protocols. The EV fraction and whole cell lysates were respectively subjected to immunoblotting.
